# Dopamine Depletion Drives Whole-Brain Oscillatory Disruptions via Cortico–Subcortical Resonance: A Multiscale Model of Parkinson’s Disease in Mice

**DOI:** 10.64898/2026.05.15.725133

**Authors:** Benedetta Gambosi, Dionysios Perdikis, Jil Meier, Alice Geminiani, Alberto Antonietti, Alberto Mazzoni, Giancarlo Ferrigno, Petra Ritter, Alessandra Pedrocchi

## Abstract

Parkinson’s disease is defined by dopaminergic neuron loss in the substantia nigra, yet its hallmark, exaggerated beta-band synchrony, pervades motor cortex, thalamus, and cerebellum, implicating network dynamics far beyond any single circuit. How focal subcortical dopamine depletion translates into brain-wide oscillatory pathology remains unresolved. We use a connectome-constrained multiscale model of the mouse brain, embedding biophysically detailed spiking networks of basal ganglia and cerebellum within whole-brain corticothalamic dynamics grounded in the Allen Mouse Brain Connectivity Atlas. We show that confining dopamine depletion exclusively to subcortical circuits is sufficient to produce widespread beta hypersynchrony (10–30 Hz), accompanied by heterogeneous theta and gamma dysregulation. Virtual loop ablations reveal that cortical and cerebellar beta amplification strictly requires intact cortico–basal ganglia–thalamic feedback; severing this loop confines beta to subcortical generators. These results support resonance within closed large-scale loops, rather than local rhythmogenesis, as the mechanism underlying distributed Parkinsonian beta pathology.

## 1 Introduction

Parkinson’s disease (PD) is increasingly recognised as a systems-level disorder whose pathology extends far beyond the basal ganglia (BG) [1]. Although the cardinal motor symptoms, rigidity, bradykinesia, and tremor, arise from dopamine loss within striatonigral circuits, structural and functional alterations are observed across thalamic, cortical, cerebellar, and other subcortical networks in patients [2–4]. Electrophysiological recordings in PD patients reveal exaggerated beta-band (approximately 13–30 Hz) synchrony across cortico–BG–thalamocortical loops in the dopamine-depleted state: subthalamic nucleus (STN) beta power correlates with rigidity and bradykinesia severity and is normalised by levodopa or deep brain stimulation (DBS) [5–7], and simultaneous magnetoencephalography (MEG) and local field potentials (LFP) recordings show that motor cortex–STN beta coherence is reduced by levodopa, with cortico-STN and cortico-muscular beta coupling in the OFF state correlating negatively with akinesia and rigidity scores [8]. During force-control tasks, stronger suppression of beta power and beta synchrony in STN and the internal Globus Pallidus (GPi) accompanies higher-force movements, further confirming that this oscillatory signature is tightly regulated by dopaminergic state [9, 10]. Cerebellar involvement is equally well-documented in humans: resting-state fMRI in early, drug-naïve PD reveals altered functional connectivity within both striatal–thalamo–cortical and cerebello–thalamo–cortical circuits, with hyperconnectivity between BG and motor cortex and between cerebellum and motor regions in tremor-dominant patients [11]. Effective connectivity Dynamic Causal Modeling (DCM) in tremor-dominant PD further traces tremor-related activity from the internal globus pallidus through motor cortex into the cerebello–thalamo–cortical loop, establishing a causal route by which BG output can recruit cerebellar circuits [12]. Together, these human data establish that pathological beta synchrony is a distributed, network-level phenomenon engaging both BG and cerebellar circuits, but its mechanistic origin remains unresolved.

Despite this rich phenomenology, the causal mechanisms underlying distributed beta synchrony cannot be resolved from human recordings alone. Clinical electrophysiology is spatially sparse and samples only a fraction of the relevant circuitry, while functional neuroimaging cannot resolve the rapid oscillatory dynamics that define the pathological state. Longitudinal tracking of disease progression is confounded by medication effects, symptom heterogeneity, and individual anatomical variability, precluding clean dissection of how dopaminergic deficits evolve into network-level pathology. Most critically, human recordings cannot distinguish whether beta rhythms emerge intrinsically within BG microcircuits and propagate outward to cortex, or whether they originate in motor cortex and are amplified by BG resonance, a distinction with direct implications for understanding and targeting the disease. Computational models that embed circuit-level perturbations within anatomically realistic whole-brain networks are therefore required to bridge these scales and probe mechanisms inaccessible to any single experimental approach.

Large-scale biophysical simulations grounded in empirical connectomes have begun to address these limitations, revealing mechanistic variables that track disease severity more closely than static functional connectivity [13]. In human-derived models, DCM of thalamocortical LFP recordings in PD demonstrates that dopamine depletion reorganises effective connectivity within cortico–BG–thalamic circuits, strengthening cortex-to-STN drive and altering indirect-pathway coupling in ways that promote pathological beta synchronisation [14, 15]; proof-of-concept work using individualised whole-brain simulations further suggests that simulation-derived parameters can augment machine-learning disease classification, though analogous approaches have not yet been established in PD [16]. In rodent-based models, a data-driven spiking BG model fitted directly to resting-state fMRI revealed Parkinsonian alterations in firing rates and effective connectivity consistent with clinical DCM findings [17]. Building on this, co-simulation architectures that embed detailed spiking BG models within The Virtual Brain (TVB)-based whole-brain dynamics have shown that local BG perturbations reshape distributed cortical activity, particularly in frontal regions, when embedded in anatomically informed large-scale models [18]. Multiscale BG–cerebellar spiking models further demonstrate that simultaneous dopamine loss in BG and cerebellum is necessary and sufficient to reproduce increased beta oscillations and motor-learning deficits [19], and virtual brain models incorporating explicit dopaminergic mechanisms capture dopamine-dependent changes in both rhythmic and aperiodic neural activity [20]. However, no existing framework integrates both BG and cerebellar spiking circuitry with a whole-brain connectome model that preserves realistic corticothalamic network topology, leaving the question of large-scale propagation unanswered.

The mechanistic origin and network-level propagation of pathological beta oscillations in PD remain actively debated. Two competing hypotheses have been formalised computationally by Pavlides and colleagues [21]: in the *intrinsic* (feedback) model, beta rhythms arise de novo within the reciprocally connected STN and the external Globus Pallidus (GPe) network and subsequently entrain cortex [22]; in the *resonance* model, motor cortex generates beta oscillations that BG circuits amplify via the hyperdirect pathway and sustain through the closed cortico–BG–thalamic loop. Experimental evidence increasingly favours the resonance view: primate recordings show that cortical beta entering the STN–GPe loop is locally enhanced and re-propagated through thalamus back to cortex [23]; DCM consistently identifies strengthened cortex-to-STN effective connectivity as the primary reorganisation following dopamine depletion [14]; and network-level analyses report that pathological beta synchrony is widely distributed and cannot be localised to any single structure, with strengthened coupling in the long cortical feedback loop from BG output nuclei via thalamus proposed as the facilitating mechanism [24]. At the same time, electrocorticography combined with STN LFPs in humans shows that tremor-related activity can transiently relieve excessive beta synchrony within the cortico–BG loop, suggesting that the two cardinal symptom clusters may involve partially distinct mechanisms [25]. Whether a closed cortico–BG–thalamic loop is a *necessary* condition for the large-scale expression of beta disruptions, as opposed to their local generation within BG microcircuits, has not been directly tested in an anatomically realistic whole-brain model.

Resolving this question requires a species in which circuit-level manipulation, multisite electrophysiology, and whole-brain structural connectivity data can all be combined at sufficient resolution. The mouse provides unmatched advantages on all three fronts. The Allen Mouse Brain Connectivity Atlas [26] delivers a mesoscale, brain-wide axonal projection dataset derived from standardised anterograde tracing experiments, constituting the most spatially complete and publicly available structural connectome for any mammalian species, providing a principled anatomical backbone for whole-brain simulation that far exceeds what is achievable from human DTI tractography. The unilateral 6-OHDA (6-Hydroxydopamine hydrobromide) mouse model reliably recapitulates the core PD electrophysiology: enhanced beta oscillations in STN and GPe local field potentials (peak 17–22 Hz under anaesthesia, 22–36 Hz in awake animals), quantifiable motor deficits on cylinder, rotarod, and open-field tasks, and partial rescue by DBS [27]. Longitudinal 6-OHDA rat recordings further show that cortical mid-gamma disruption (41–45 Hz) emerges within hours of lesion induction, whereas pathological beta synchrony in motor cortex and BG builds gradually over days [28], establishing a well-characterised temporal progression suitable for model validation. Region-specific dopaminergic alterations along BG–cortico–cerebellar pathways have been characterised at single-cell resolution in mice [29, 30], providing the parametric constraints needed to build biologically realistic spiking models that are currently unavailable from human tissue. Our group’s prior BG–cerebellar multiscale model [19] confirmed in this rodent framework that simultaneous dopamine loss in BG and cerebellum best accounts for thalamocortical beta abnormalities, but represented all cortical dynamics as a single lumped node, precluding investigation of how dopaminergic perturbations propagate through large-scale corticothalamic networks. We therefore extend this preclinical foundation into an anatomically realistic, Allen-connectome-based whole-brain framework.

To address this limitation, we integrate the Allen Mouse Brain Atlas connectome into the TVB framework [31, 32], using a TVB–spiking co-simulation architecture comprising corticothalamic neural mass nodes coupled to detailed spiking BG and cerebellar circuits [18, 33]. Dopamine depletion is implemented *exclusively* within the BG and cerebellar spiking neural networks (SNNs), while cortical and thalamic dynamics remain governed by physiological neural mass models, a design that isolates the propagation of local dopaminergic deficits through the structural connectome from any assumed change in cortical excitability. Spiking networks are essential for the BG and cerebellar components because dopamine-dependent microcircuit mechanisms, including altered BG cell excitability and cerebellar apoptosis, cannot be faithfully represented in mean-field approximations without sacrificing biologically realistic firing patterns [20]. Crucially, targeted *in silico* connectivity perturbations, selectively severing or scaling specific inter-regional pathways, enable causal dissection of loop contributions that is impossible in vivo.

Using this framework, we specifically ask: (i) How does dopamine depletion confined to BG and cerebellar microcircuits alter oscillatory dynamics across the whole mouse brain? (ii) Is widespread cortical beta synchrony generated locally within BG circuits, or does it require resonance within a closed cortico–BG–thalamic loop? Through systematic whole-brain simulations and targeted loop perturbations, we show that distributed beta-band hypersynchrony arises from the interaction between locally altered BG and cerebellar circuits and recurrent cortico-subcortical loops, supporting a resonance-based mechanism for PD rhythmopathies and demonstrating that anatomically realistic large-scale connectivity is a necessary condition for the network-wide expression of pathological beta synchrony.

## 2 Results

### 2.1 A multiscale whole-brain model linking dopaminergic microcircuits to network dynamics

We first integrated dopamine-sensitive spiking circuits of the BG and cerebellum into a mouse-specific whole-brain model that integrates anatomically resolved corticothalamic population dynamics (derived from the Wilson-Cowan model [33, 34]). The large-scale backbone consisted of a whole-brain network with connections derived from the Allen Mouse Brain atlas [26], and corticothalamic nodes [26], comprising cortical, thalamic, basal ganglia, cerebellar, and additional subcortical nodes (Fig. 1). Each region, apart from BG and cerebellum nodes, was implemented as a four-population corticothalamic neural mass model [35], supporting oscillatory dynamics in the alpha–beta range and capturing canonical cortico–thalamic feedback loops (see Methods Subsection 5.2).

**Fig. 1:**
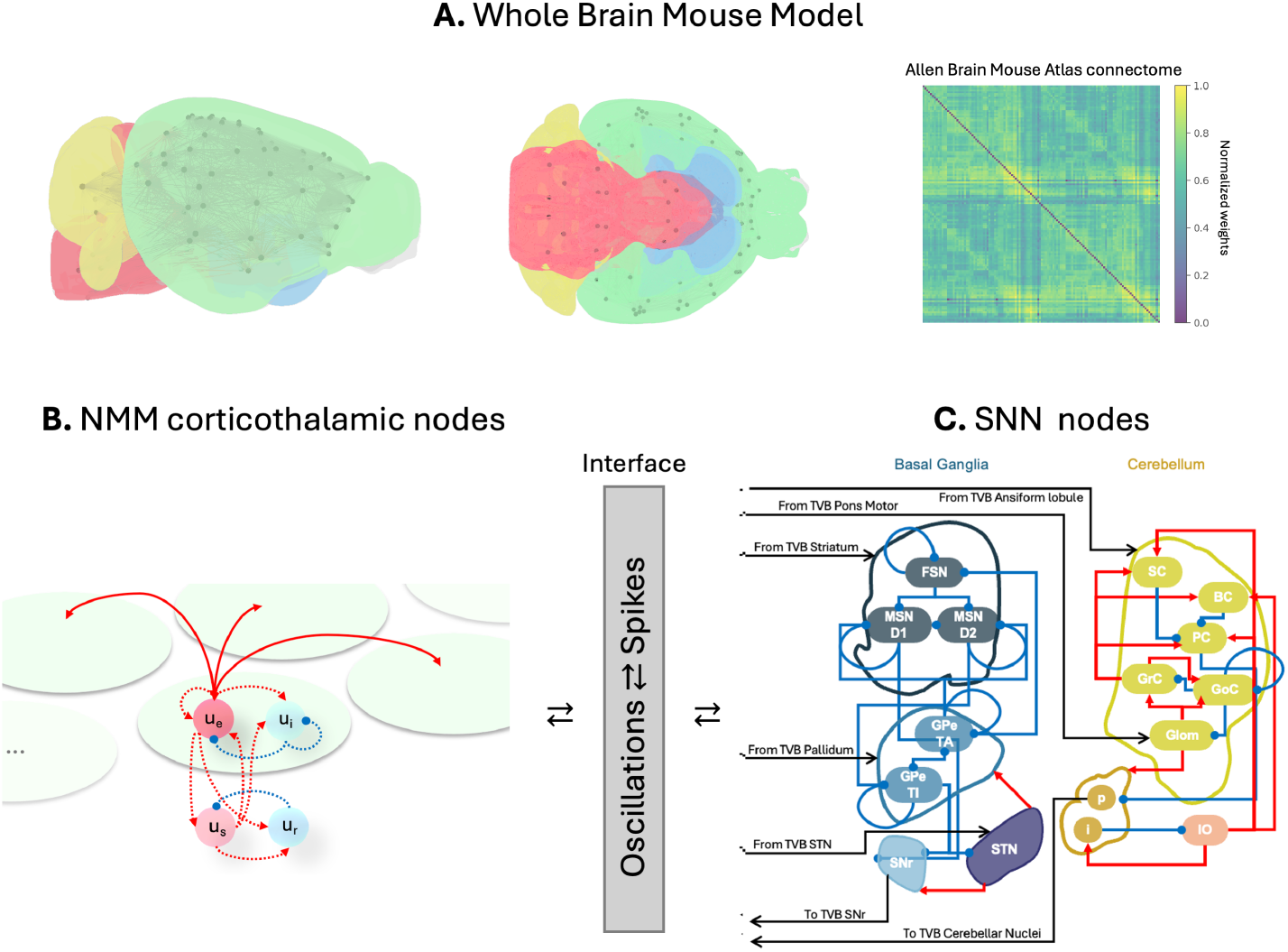
Multiscale whole-brain model of dopaminergic perturbation in the mouse. A. Whole-brain network derived from the Allen Mouse Brain Atlas [26], comprising bilateral cortical (green), subcortical (cyan), cerebellar (yellow), and brainstem (magenta) regions coupled via empirical structural connectivity. **B. Corticothalamic neural mass model (NMM) node structure**: each oval represents a distinct brain region (ROI) hosting its own instance of the NMM. Reddish spheres denote excitatory populations (*u_e_*, *u_i_*) and blue spheres inhibitory populations (*u_s_*, *u_r_*), coupled through local delayed feedback loops (dotted arrows). Red arrows between ovals represent long-range anatomical projections connecting cortical nodes across regions, scaled by structural connectome weights. **C. Spiking neural network (SNN) modules** for basal ganglia (in shades of blue) and cerebellum (in shades of yellow), incorporating dopamine-sensitive microcircuit mechanisms. Red arrows indicate excitatory connections, blue arrows inhibitory ones. Arrows ending to area outline connect to all inside population. Bidirectional interfaces convert TVB firing rates into Poisson spike trains and return spiking activity as low-pass filtered firing-rate signals, enabling coherent co-simulation across scales. Black arrows in panel **C** indicate connections with the TVB nodes (details in Table 1).

**Table 1:**
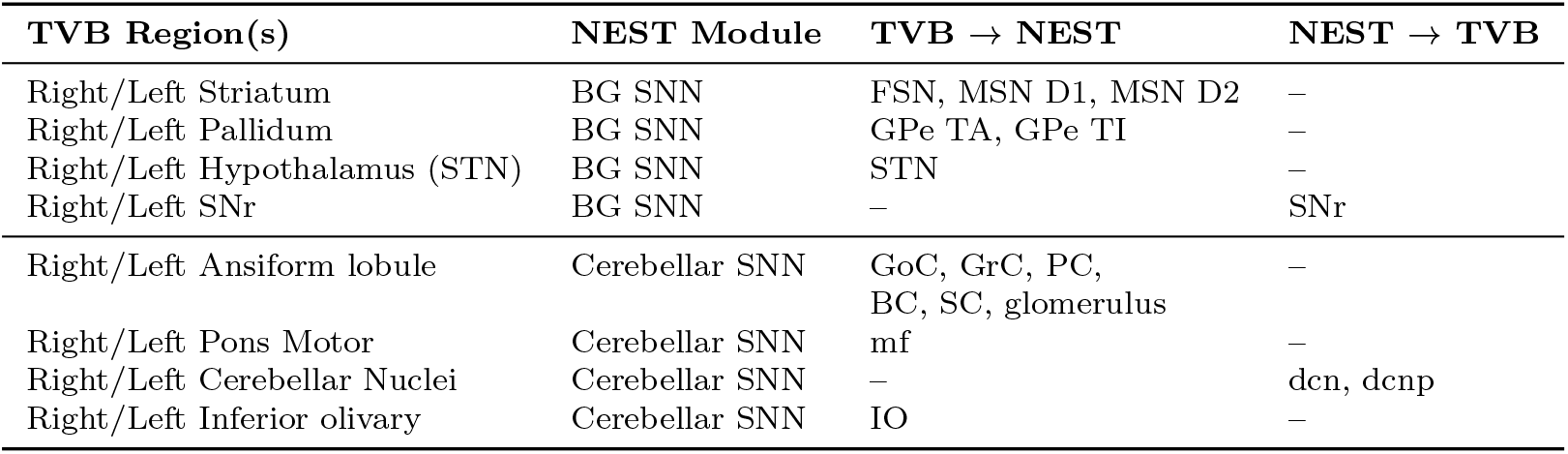
TVB-NEST region-to-population mapping for co-simulation interfaces.

To incorporate circuit-level dopaminergic mechanisms, we leveraged the multiscale framework from a previous work by our team [19] and embedded two detailed spiking microcircuits: a BG model adapted from Lindahl and Kotaleski [36] and a cerebellar olivocerebellar circuit adapted from Geminiani et al. [37]. These models include experimentally grounded mechanisms for dopamine depletion (*ξ*), enabling us to manipulate BG and cerebellar dopamine levels (see Methods Subsection 5.4).

The spiking microcircuits and the whole-brain model were coupled through a multiscale co-simulation interface [18, 33], allowing BG and cerebellar activity to interact bidirectionally with detailed large-scale corticothalamic dynamics (see Methods Subsection 5.3). This architecture preserves the intrinsic behaviour of the microcircuits while enabling dopamine-dependent perturbations to propagate through the structural connectome and influence global network rhythms. This multiscale integration allowed us to examine how focal dopaminergic deficits reshape distributed oscillatory patterns across the mouse brain.

### 2.2 Dopaminergic perturbation drives distributed spectral dysregulation

We simulated a physiological baseline, previously validated against rodent LFP spectra in the corticothalamic neural mass model (NMM) framework (Griffiths et al., [35]; adapted for mouse [33]) and single-unit recordings in standalone BG/cerebellar circuits [19, 36, 37].

The Parkinsonian condition implemented dopaminergic depletion by increasing subthalamic nucleus (STN) and external Globus Pallidum (GPe) excitability, reducing internal Globus Pallidus (GPi) inhibition, and increasing deep cerebellar nuclei (DCN) output through Purkinje cells (PC) apoptosis [19, 38]. Minor adaptations to the multiscale TVB-SNN integration preserved these validated dynamics, confirmed by consistency checks (Figure S1 in Supplementary information).

We performed two complementary analyses. First, to characterise how pathology severity shapes the canonical Parkinsonian rhythmopathy, we focused on beta-band activity across increasing levels of dopamine depletion. Second, for the most severe condition, we examined the full spectral profile by estimating power across delta (0.5–4 Hz), theta (4–10 Hz), beta (10–30 Hz), and gamma (30–100 Hz) bands using the continuous wavelet transform (see Methods Subsection 5.6).

Relative changes were computed against baseline, and statistical significance was assessed using Wilcoxon–Mann–Whitney tests with FDR correction. In what follows, we report only those macro-domains exhibiting at least one significant beta-band change across pathology levels.

To situate these spectral alterations within the model’s structural organisation, we grouped regions into functional macro-domains (see Methods Subsection 5.7) (Fig. 2). The structural connectome displays both within-hemisphere and cross-hemispheric interactions, with the majority of connections converging on subcortical structures. The thalamus and BG acts as central hub, projecting broadly to virtually almost all other regions across both hemispheres. Strong intra-hemispheric pathways link the thalamus and basal ganglia, while the thalamus and pons form prominent inter-hemispheric connections with their homotopic counterparts. The STN, midbrain, pons and cerebellum participate in tightly interconnected subcortical loops. This anatomical organisation provides a mechanistic substrate for the observed progression of beta dysregulation, from early subcortical amplification centred on the basal ganglia–thalamus–STN axis to later involvement of interconnected cortical and cerebellar systems.

**Fig. 2:**
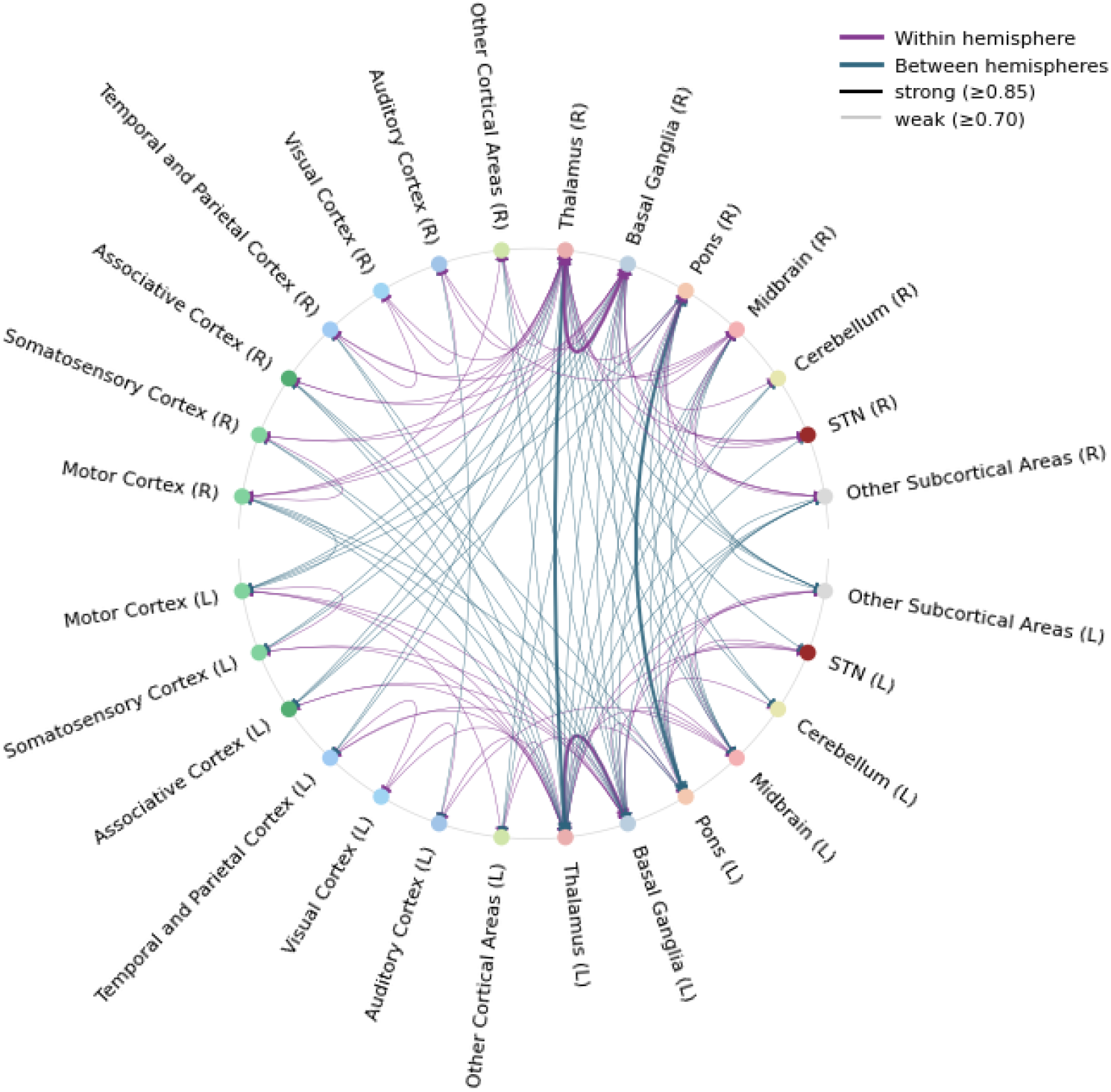
Connection scheme of functional macro-domains used to interpret and summarise spectral changes across the mouse brain. Red edges denote within-hemisphere connections and blue edges between-hemisphere (interhemispheric) connections. Line thickness encodes connection strength: thick lines indicate strong projections (≥ 0.85) and thin lines weaker projections (≥ 0.7), derived from the Allen Mouse Brain Atlas structural connectome (connection weights < 0.70 are not shown for readability). Macro-domains colored according to Allen Mouse Brain Atlas ontology. Notable dense interhemispheric coupling is visible between bilateral pons and thalamus nodes, reflecting the strong cross-hemispheric anatomical projections of these structures, and within-hemispheric connection between thalamus and BG.

#### (i) Circuit-level dopamine perturbation reshapes beta dynamics

Dopaminergic loss produced a progressive, anatomically structured increase in beta-band power across the network (Fig. 3). Subcortical structures emerged as primary generators with pronounced, monotonic amplification, while cortical involvement was delayed and strictly domain-specific.

**Fig. 3:**
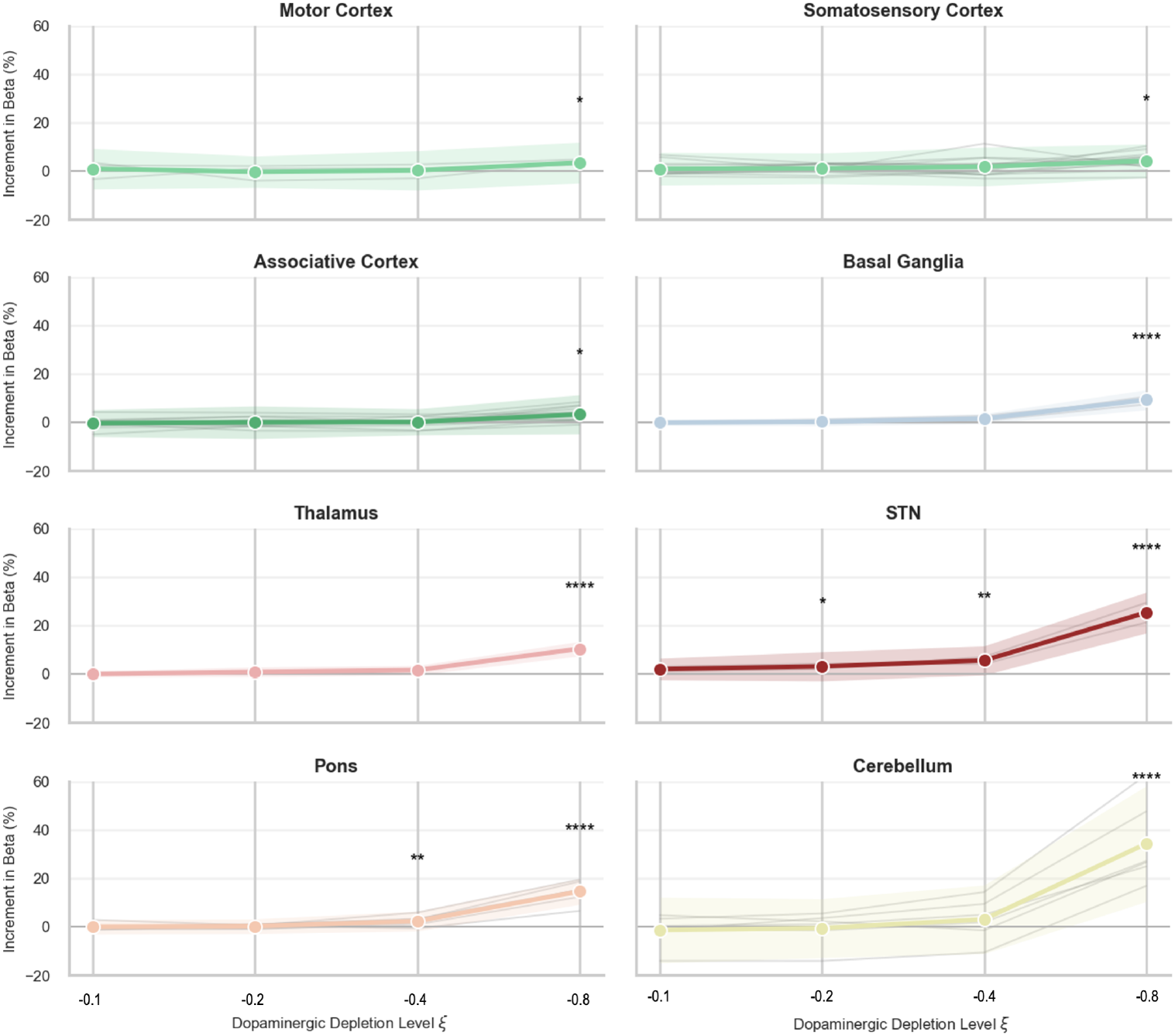
Progressive dopaminergic depletion induces region-specific increases in beta-band power across the mouse brain. For each macro-domain, the percentage change in beta power relative to physiological baseline is shown across four levels of dopamine depletion (*ξ* = −0.1, −0.2, −0.4, −0.8). Grey lines represent individual regions within each macrodomain; bold curves denote group averages with shaded 95% confidence intervals. Among subcortical structures, STN and Pons exhibit earlier and steeper monotonic increases, whereas the basal ganglia, thalamus, cerebellum and cortical territories (Motor, Somatosensory, Associative) display significant beta enhancement only at the most severe depletion level. Asterisks indicate FDR-corrected significance levels (* *p <* 0.05; ** *p <* 0.01; *** *p <* 0.001).

**Subcortical effects:** SNN-embedded *STN* showed the earliest and a steep pathology-dependent rise driven by local dopaminergic effects. The *Pons* and *thalamus* exhibited reliable beta elevation despite lacking dopamine modulation, acting as a key relay. The *cerebellum* displayed substantial late-onset increases, despite DCN suppression by Purkinje apoptosis. These patterns reveal subcortical nuclei as the origin of pathological beta, progressively recruiting connected structures.

**Cortical effects:** Among cortical domains, significant beta augmentation emerged only at peak severity in *motor*, *somatosensory*, and *associative* cortices. Temporal/parietal, visual, auditory, and other cortical areas remained unaffected, underscoring anatomical selectivity despite shared Wilson-Cowan dynamics.

This hierarchical gradient (STN > subcortical ≥ cortical) demonstrates propagation governed by connectome topology (derived from the Allen Mouse Brain Atlas [26]).Uniform model equations across nodes produced domain-specific effects, confirming connectome-driven resonance as the organizing principle.

#### (ii) Advanced pathology disrupts cortical and subcortical frequency structure

Robust beta-band alterations prompted us to extend the analysis to other frequency bands in the most severe condition (*ξ* = −0.8).

*Subcortical spectral alterations under dopaminergic depletion.* Dopaminergic loss produced pronounced and spatially coherent spectral abnormalities across subcortical systems, with the strongest and most consistent effects in the beta band, together with region-specific changes in theta and gamma activity (Fig. 4). Such beta-band enhancement is consistent with experimental and clinical evidence of pathological synchronization in Parkinsonian cortico-BG circuits [21, 39–42]. Only regions exhibiting at least one significant change are described below.

- The **basal ganglia** showed a significant increase in beta power and a decrease in theta and gamma power. Increased beta activity in basal ganglia circuits is well supported by experimental studies in Parkinsonian rodent models and patients [21, 39, 40, 42, 43]. Our model further predicts that this beta enhancement arises from the interaction between local dopaminergic microcircuit changes and network-level reverberation within BG loops (see also Subsection 2.3).
- **Thalamic nuclei** displayed significant increases in beta power and decreases in the other bands. The involvement of thalamic relay pathways in Parkinsonian beta synchronization is consistent with experimental and modeling work on cortico-thalamo-BG interactions [14, 15, 42]. A model-specific prediction is that thalamic beta elevation should scale with the strength of subcortical feedback and could therefore be tested by selectively perturbing thalamic relay activity.
- The **STN** exhibited a robust increase in beta power and a decrease in theta power, with beta reaching the highest levels observed across the subcortical groups. STN beta activity is one of the most robust experimental signatures of Parkinsonism and has been repeatedly linked to dopaminergic depletion and motor impairment [7, 39–41, 44]. This result suggests that this STN beta amplification is a necessary hub-level mechanism for propagating pathological rhythms across the subcortical network.
- **Pontine regions** showed significant increases in beta, and a decrease in theta and gamma power. While the beta elevation suggests integration into the broader pathological oscillatory network, the concurrent gamma decrease points to altered local circuitry within brainstem–cerebellar pathways. These findings indicate that brainstem structures could actively participate in the distributed frequency reorganisation induced by dopamine loss.
- **Cerebellar regions** demonstrated marked increases in beta power. This modulation indicates that cerebellar circuits do not remain spectrally independent of basal ganglia–thalamic dynamics but instead undergo frequency-specific entrainment [45]. As previously found [19], the cerebellum actively contributes to, and not only reflects, the broader network-level reconfiguration associated with Parkinsonian pathology.

**Fig. 4:**
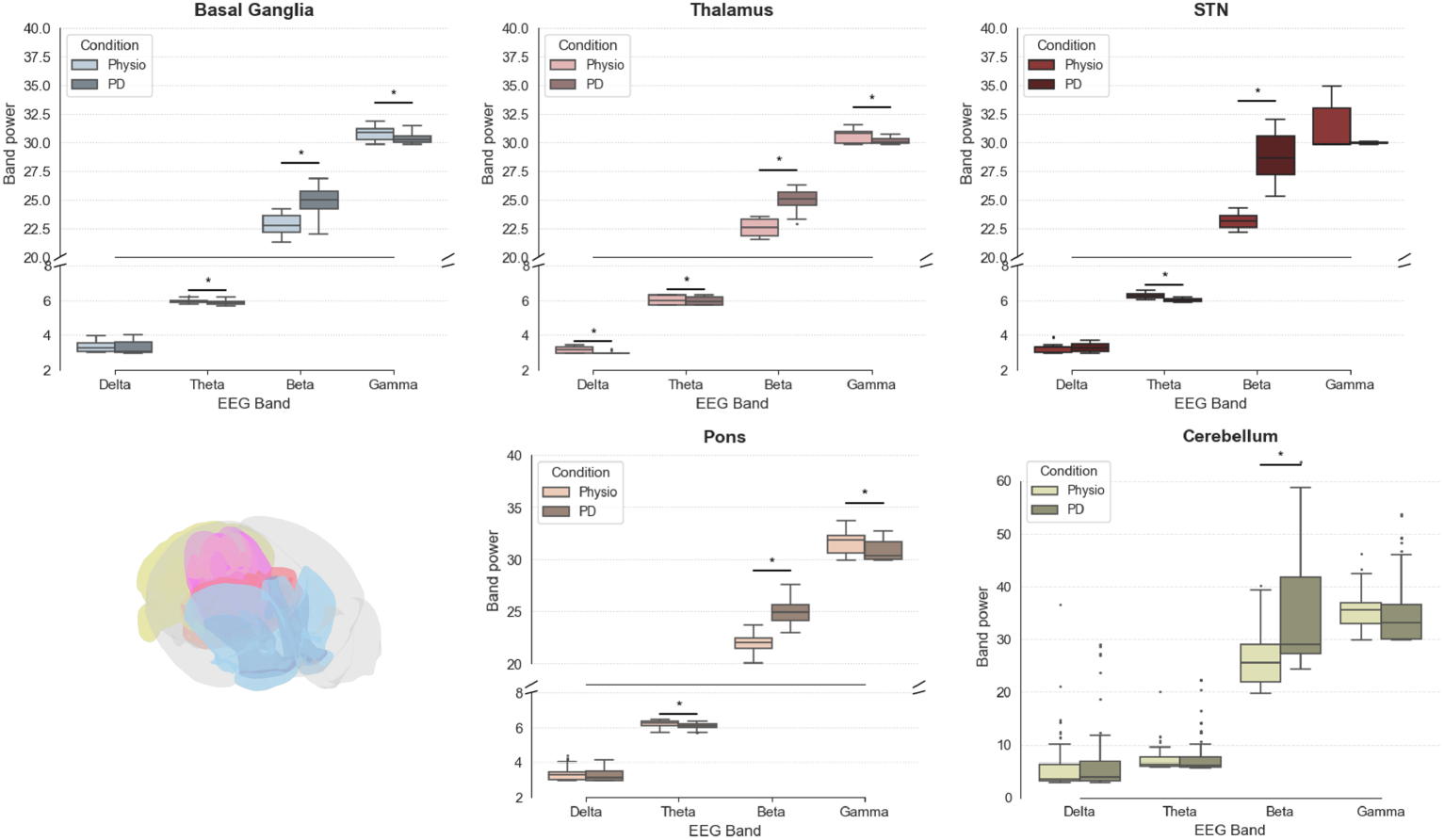
PSD change per band across subcortical macro-domains (basal ganglia, thalamus, STN, cerebellum, pons, midbrain). Beta-band increases dominate in BG, thalamus, and cerebellum, whereas gamma suppression is prominent in midbrain structures. Boxes represent distributions across regions and physio-pathological conditions; stars indicate significant differences: Wilcoxon–Mann–Whitney tests with FDR correction (∗ : *p* < 0.05).

Together, these results reveal a consistent gradient of subcortical involvement, with beta-band hypersynchrony emerging as a network-wide hallmark of dopaminergic depletion and theta/gamma alterations reflecting region-specific circuit adaptations. The pattern supports a view of Parkinsonian oscillations as a distributed subcortical phenomenon rather than one confined to classical basal ganglia loops.

*Cortical spectral alterations under dopaminergic depletion.* Cortical macro-domains exhibited frequency-specific modulations in response to dopaminergic perturbation (Fig. 5). Across all three domains displaying significant changes, alterations were confined to the beta and gamma bands, with delta and theta activity remaining largely unaffected. This pattern is consistent with the broader literature on Parkinsonian cortex, which reports prominent beta-band abnormalities [42] and, in several contexts, concomitant changes in gamma activity [46–48].

- The **motor cortex** showed a significant increase in beta-band power in the Parkinsonian condition, whereas gamma-band activity remained unchanged. Increased cortical beta activity is consistent with experimental and clinical reports of Parkinsonian motor-cortex involvement and with the view that beta hypersynchrony extends beyond the basal ganglia into cortical motor circuits [42, 48, 49].
- The **somatosensory cortex** exhibited a robust beta-band increase as well as a smaller but significant elevation in gamma power. Experimental work supports the involvement of sensorimotor and somatosensory regions in Parkinsonian network dynamics, including altered beta-related coupling and impaired sensorimotor processing [42, 50]. The additional gamma increase in our simulations is a model-specific prediction, suggesting that somatosensory cortex may be more strongly recruited than beta-only descriptions would imply, potentially reflecting impaired sensory integration and reduced adaptability of sensorimotor loops under dopaminergic loss [51].
- The **associative cortex** demonstrated significant increases in both beta and gamma power. Cortical involvement outside primary motor areas is consistent with the broader literature on Parkinsonian network reorganization and cognitive-motor dysfunction [42, 49, 50]. Such alterations may correspond to executive and cognitive inflexibility frequently observed in Parkinsonian states [52].

**Fig. 5:**
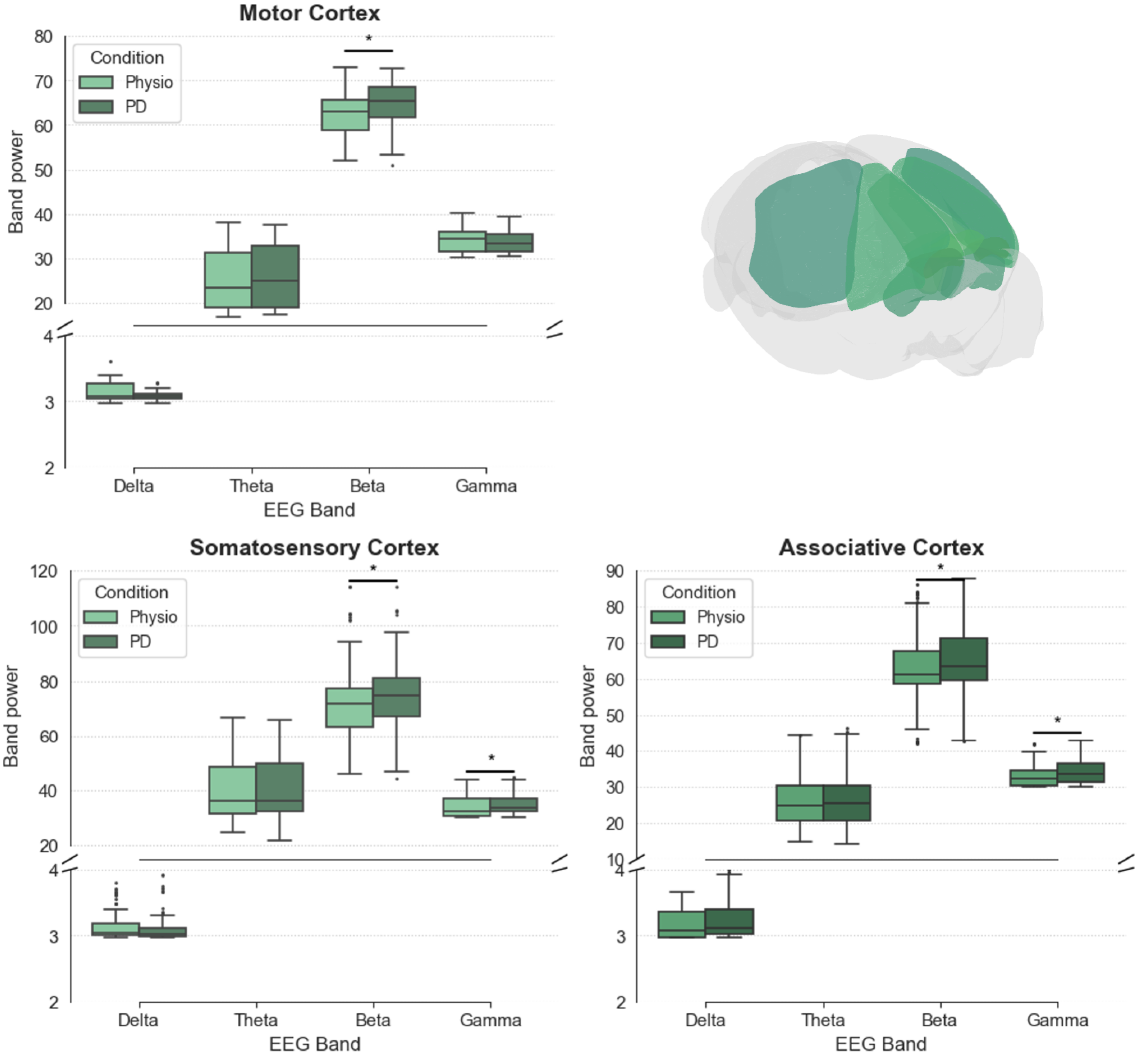
PSD change per band across cortical macro-domains (motor, somatosensory, associative). Boxes represent distributions across regions and physio-pathological conditions; asterisks indicate significant differences: Wilcoxon–Mann–Whitney tests with FDR correction (∗ : *p* < 0.05).

Overall, these cortical findings reveal a graded and domain-specific pattern: beta-band hypersynchrony emerges across motor, somatosensory, and associative cortices, accompanied by gamma-band increases in the latter two domains. Although sharing same dynamical equations, cortical domains differ mechanistically via connectome: motor cortex receives strongest hyperdirect (cortex-STN) and indirect (cortex-GPe) feedback from BG SNNs, yielding selective beta; somatosensory amplifies both via sensorimotor thalamus loops plus gamma from pontine-cerebellar redistribution; associative shows dual beta/gamma due to diffuse frontal projections diluting SNN signals but preserving E/I imbalance. These are not generic “sensorimotor/executive” effects but emerge from the Allen connectome topology [26] and SNN-to-TVB rate mapping. These results support the view that cortical involvement in Parkinsonian dynamics is secondary yet progressively recruited as pathological subcortical oscillations propagate through connected networks.

### 2.3 In-silico feedback ablation experiments to probe the origin and propagation of beta synchrony

To dissect the respective contributions of intrinsic BG rhythmogenesis and cortico–subcortical resonance, we performed a targeted connectivity manipulation in which all feedback projections from cortex and thalamus to the BG were selectively removed, while preserving intra–BG connectivity and all outgoing outputs (see Subsection 5.8). This perturbation allowed us to isolate the intrinsic rhythmogenic capacity of BG circuits from the re-entrant dynamics of the larger network.

In the in-silico feedback ablation experiment, beta power remained significantly elevated only in BG and Pons (Fig. 6). These results show that a part of subcortical circuits can sustain beta oscillations autonomously under dopaminergic depletion, consistent with intrinsic rhythmogenic mechanisms within the BG.

**Fig. 6:**
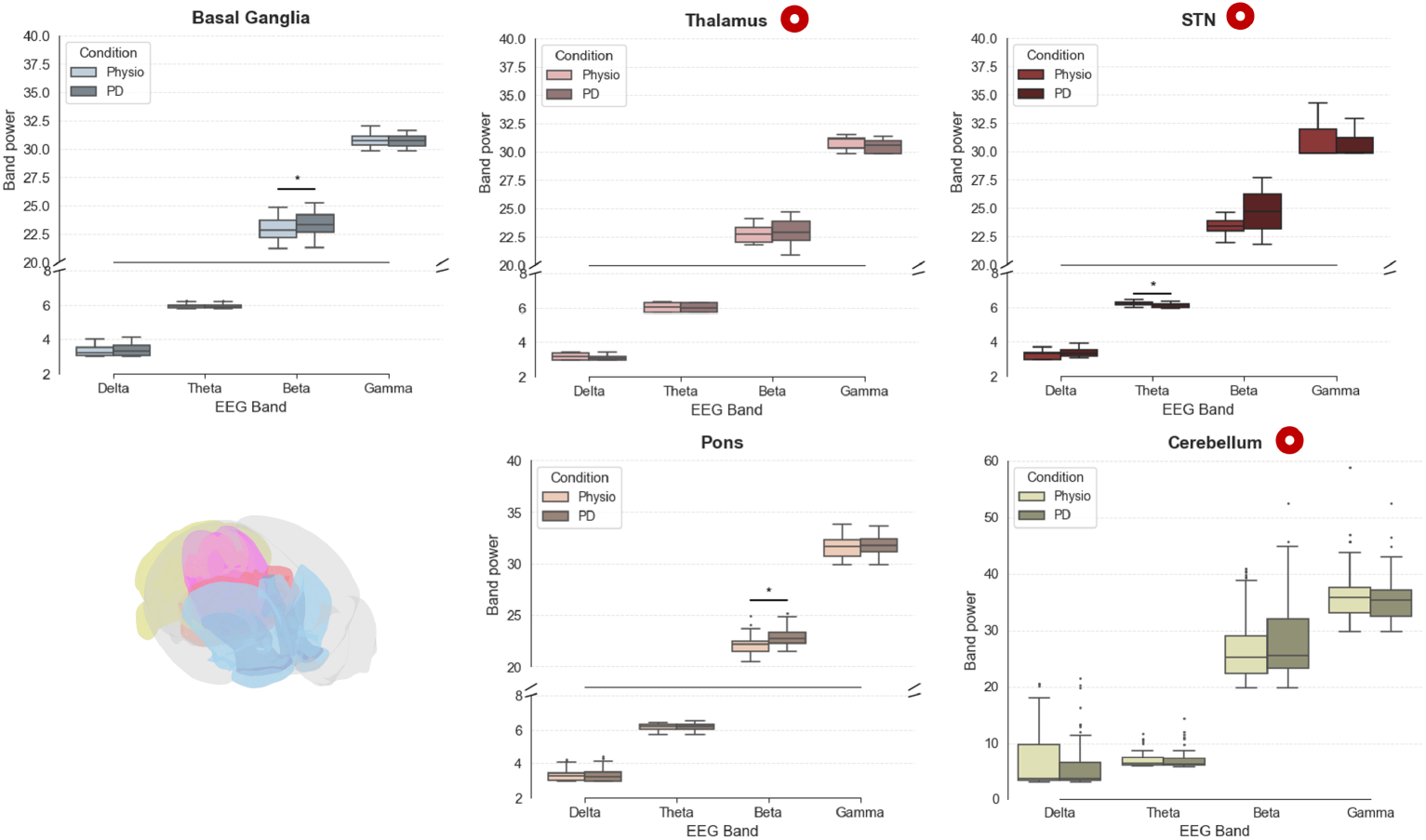
In-silico feedback ablation experiment results in subcortical macro-domains. Beta power remains elevated in BG and Pons in the in-silico feedback ablation experiment, indicating that local oscillatory generation persists despite loss of re-entrant loops. Red markers highlight macro-domains that lose significant beta enhancement when cortico–BG feedback is removed. Asterisks indicate significant differences: Wilcoxon–Mann–Whitney tests with FDR correction (∗ : *p <* 0.05)

Disconnecting cortex/thalamus → BG feedback while preserving intra-BG connectivity and efferents isolated SNN rhythmogenesis from re-entrant amplification. Beta power remained elevated only in BG/Pons via local dopaminergic loops; thalamus retained partial relay via SNr; but cortex and cerebellum collapsed without cortico-BG feedback (Fig. 6).

Cerebellum showed minimal direct change despite local *ξ* because its pathological DCN output requires loop-mediated mossy-fiber entrainment from pontine beta. Without feedback, DCN spikes saturate locally without global propagation. Cortical macro-domains (motor, somatosensory, associative) lost all PD-specific beta/gamma effects, confirming closed corticothalamic loops as necessary for network-wide expression (Fig. 7).

**Fig. 7:**
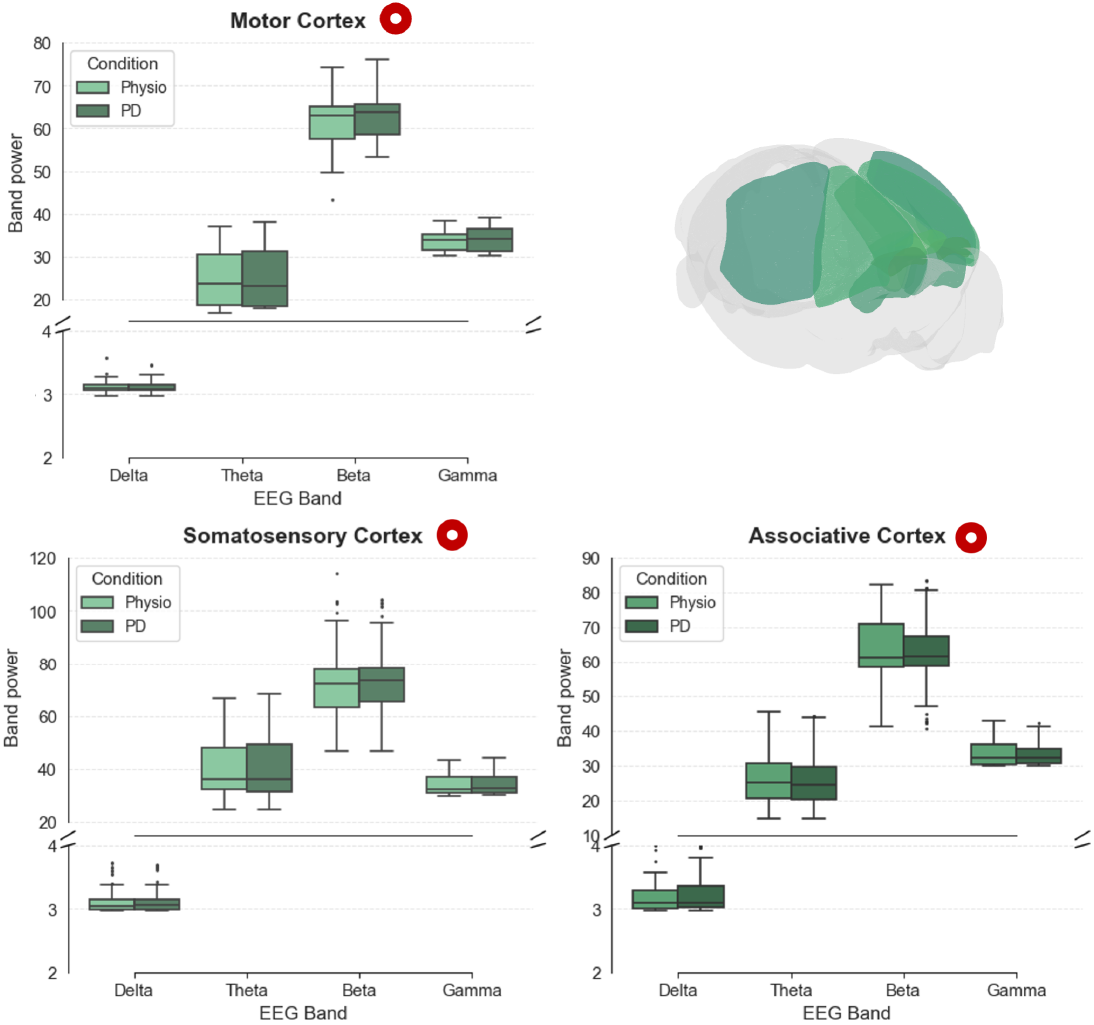
In-silico feedback ablation experiment results in cortical macro-domains. Comparison of beta power changes between full Parkinsonian configuration and in-silico feedback ablation experiment. Red markers highlight macro-domains that lose significant beta enhancement when cortico–BG feedback is removed. Asterisks indicate significant differences: Wilcoxon–Mann–Whitney tests with FDR correction (∗ : *p <* 0.05)

Taken together, these findings indicate that while dopaminergic loss renders BG circuits capable of generating beta oscillations locally, the widespread propagation and amplification of these oscillations across cortex, brainstem, and cerebellum critically depends on the integrity of closed cortico–BG–thalamic loops. Disrupting feedback effectively confines pathological beta synchrony to subcortical nodes and prevents its large-scale expression. This behaviour is consistent with resonance-based accounts of Parkinsonian dynamics, in which re-entrant coupling between cortex and basal ganglia stabilises and broadcasts pathological beta activity [21, 53].

## 3 Discussion

### 3.1 Main findings

The simulations directly address the two questions posed in the Introduction (Section 1). First, dopaminergic perturbation confined to BG and cerebellar spiking circuits is sufficient to induce anatomically precise, frequency- and region-specific alterations in whole-brain dynamics, arising naturally from the interaction between local microcircuit changes and the Allen Mouse Brain Atlas-based large-scale network structure [26], without parameter tuning to enforce pathological rhythms and without any assumed change in cortical excitability. The resulting patterns reproduce hallmark electrophysiological features of Parkinsonian activity, including widespread beta-band augmentation and selective gamma- and theta-band reorganisation, highlighting how focal circuit-level deficits propagate through the mouse connectome to shape global neural states. Second, and more mechanistically, the feedback ablation experiment demonstrates that this network-wide expression of beta hypersynchrony requires a closed cortico-BG-thalamic loop: local STN-GPe beta persists after loop severance, but synchrony collapses across cortex and cerebellum. Together, these findings directly support the resonance hypothesis formalised by Pavlides and colleagues [21] and contradict a purely intrinsic origin of large-scale beta pathology.

A central finding is the hierarchical progression of beta abnormalities. The strongest and earliest enhancements occurred in STN, basal ganglia, and thalamus, consistent with experimental and computational evidence identifying the STN–GPe loop as a primary source of pathological beta oscillations [54, 55]. Beta augmentation also emerged in cerebellar nuclei and pontine pathways, aligning with rodent studies showing cerebellar beta entrainment after dopamine depletion [1, 19, 56]. Only at severe depletion levels did motor, somatosensory, and associative cortical domains show modest, but significant, beta increases, echoing mouse LFP and calcium-imaging data in which cortical beta reflects propagated, rather than intrinsic, dysregulation [57], and human MEG-LFP findings in which motor cortex–STN beta coherence is reduced by levodopa and correlates with akinesia and rigidity in the OFF state [8]. This progression reinforces the notion that pathological beta originates subcortically and spreads through structured corticothalamic loops rather than emerging uniformly across the cortex.

These hierarchical effects also address how cerebellar manipulation (*ξ* in DCN/PC) first influences BG before returning to cerebellum. Both SNNs receive identical TVB coupling *C_i_*(*t*) = *G _j_ w_ij_S*(*E_j_*(*t − τ_ij_*)), but BG drives thalamus via SNr output, which relays beta-enhanced *c*^th^ back to cerebellum through pontine mossy-fiber scaling. Thus, the sequence is: cerebellar *ξ →* TVB pons → thalamus → BG SNN → SNr → thalamus → TVB cortex → pons → cerebellar SNN. Removing corticothalamic-BG feedback (*γ*^TVB→BG^ = 0) breaks this loop, confining cerebellar beta to local *ξ*-effects without global propagation. SNN input to TVB acts as an additive coupling term in the corticothalamic model equations (Equation 2-Equation 5), injecting beta-enriched spectra that shift the excitatory drive *Gc_j_^sb^* without inducing bifurcations: pathology emerges from resonant amplification within the connectome topology rather than local instabilities.

Oscillatory alterations beyond beta further point to frequency-specific circuit imbalance. Gamma suppression in BG output nuclei and superior colliculus parallels midbrain deficits in fast-timescale coordination [48], whereas localized gamma increases in frontal regions may reflect compensatory recruitment of parallel pathways [58]. The model also reproduces the temporal ordering reported in longitudinal 6-OHDA rat recordings [28], where mid-gamma disruption emerges acutely and cortical beta synchrony builds gradually: in our simulations, gamma suppression in BG output nuclei and superior colliculus is present at intermediate depletion levels, whereas cortical beta increases appear only at severe depletion and depend on the progressive recruitment of corticothalamic feedback. Theta and delta rhythms were consistently reduced across all regions where significant effects were observed, indicating a broad suppression of slower oscillatory activity under Parkinsonian conditions rather than a regionally selective reorganization. Together with the band- and region-specific pattern of beta changes, the model supports a view of PD as a disorder of dynamic imbalance, where distinct oscillatory channels are differentially shaped by the specific topology of BG–thalamic–cortical–cerebellar loops rather than a uniform global shift toward slower rhythms.

The in-silico feedback ablation experiment provides a direct test of the two mechanistic hypotheses formalised by Pavlides and colleagues [21]. When cortico–thalamic feedback onto BG was removed, STN–GPe beta oscillations remained robust, confirming that the BG microcircuit can sustain intrinsic beta under dopamine depletion, consistent with the intrinsic (feedback) model [21, 55]. However, global beta synchrony collapsed across cortex and cerebellum, demonstrating that network-wide hypersynchrony is not an intrinsic BG property but instead requires re-entrant resonance within the closed cortico–BG–thalamic loop [53, 59]. This reconciles long-standing views in the PD literature: while intrinsic loop dynamics generate beta autonomously within the BG, but the large-scale expression of pathological synchrony, the electrophysiological hallmark that correlates with rigidity and bradykinesia severity in patients [5, 6], is a resonance phenomenon requiring the full corticothalamic network topology. Our results therefore support the conclusion, suggested by DCM analyses [14] and primate LFP work [23], that strengthened cortex-to-STN drive and long-loop cortico-BG-thalamic coupling are the primary reorganisation following dopamine depletion, rather than a change in intrinsic BG oscillatory propensity. The emergence of pathological beta is therefore best understood not as a property of any single nucleus, but as a property of a distributed resonant ensemble formed by recurrent excitatory–inhibitory loops spanning cortex, thalamus, BG, and cerebellum.

Cerebellar and brainstem structures play a particularly active role. Coherent beta enhancement in cerebellum, thalamus, and pons indicates that these regions do not simply reflect BG dysfunction but instead participate directly in the pathological loop. This is consistent with resting-state fMRI evidence in early, drug-naïve PD showing hyperconnectivity within cerebello-thalamo-cortical circuits alongside BG-motor cortex hyperconnectivity [11], and with DCM tracing tremor-related activity from GPi through motor cortex into the cerebellar loop [12]. Our results provide a mechanistic substrate for these empirical observations: BG output via SNr drives thalamus, which relays beta-enriched signals to cerebellum through pontine mossy-fibre pathways, establishing the causal route hypothesised from human DCM data. This also aligns with tract-tracing work demonstrating cerebello–thalamo–striatal pathways [60] and multiscale modelling studies showing cerebellar contributions to Parkinsonian beta and learning deficits [1, 19].

### 3.2 Position within existing multiscale modelling frameworks

Methodologically, this work extends previous multiscale simulations [18, 33, 61] by embedding biophysically grounded BG and cerebellar spiking networks into a high-resolution mouse whole-brain model. Crucially, this co-simulation architecture fills the gap identified in the Section 1: by embedding BG and cerebellar spiking circuitry within an Allen-connectome-based whole-brain model with realistic corticothalamic topology, it becomes possible to track how cerebellar dopaminergic perturbations propagate globally, a question that was unanswerable in our prior framework [19], which represented all cortical dynamics as a single lumped node. This integration enables the detection of oscillatory disruptions in underrepresented structures such as the cerebellum, pons, and midbrain. The co-simulation interface preserves intrinsic SNN dynamics while enabling stable coupling with TVB-based cortical and thalamic populations, allowing pathological signatures to arise from the interplay of micro- and macro-scale dynamics. This highlights the explanatory power of multiscale data-driven models for conducting in-silico experiments to test hypotheses that would be challenging to evaluate experimentally.

### 3.3 Limitations

While being adequate to answer the questions in this paper, several limitations of the model remain. Dopaminergic effects are implemented through static parameter modulation rather than explicit modelling of SNc neurons, receptor kinetics, or phasic dopamine dynamics [62, 63]. The lack of SNc projections limits the model’s ability to capture progressive neurodegeneration or reward-dependent modulation [64]. Validation against invasive recordings in dopamine-depleted mice, such as LFP, widefield calcium imaging, or functional ultrasound, will be necessary to establish quantitative predictive accuracy. Finally, structural connectivity is static, preventing the modelling of axonal degeneration or compensatory anatomical reorganisation [65, 66].

### 3.4 Future work

Future developments may incorporate explicit dopaminergic populations, receptor-specific mechanisms, and structural plasticity rules, as well as Simulation-Based Inference [67] to estimate parameters for latent disease states directly from mouse recordings, with validation against real experimental data to assess model predictions and refine parameter estimates. The present framework also provides a platform for testing targeted interventions such as deep brain stimulation [68, 69], optogenetic stimulation [70, 71], and cerebellar modulation [72, 73]. More broadly, our results underscore how multiscale computational models can reveal how focal cellular changes scale into distributed network dysfunction, and how restoring balance in specific loops may disrupt pathological resonance across the entire brain.

## 4 Conclusion

We developed an anatomically resolved multiscale model of the mouse brain that integrates connectome-based corticothalamic dynamics with dopamine-sensitive basal ganglia and cerebellar spiking circuits. By confining dopamine depletion to subcortical and cerebellar modules, we showed that local microcircuit perturbations are sufficient to generate widespread beta-band hypersynchrony across cortex, thalamus, and cerebellum, alongside nuanced changes in theta and gamma activity. A targeted in-silico disconnection experiment revealed that this large-scale beta propagation critically depends on closed-loop cortico–BG–thalamic resonance: subcortical beta persists without feedback, but cortical and cerebellar beta waves collapse.

These findings support a view of Parkinson’s disease as a disorder of distributed network coordination emerging from the interaction of locally altered BG and cerebellar circuits with the structural connectome. In addition, the presented framework provides a mechanistic *in silico* platform for testing hypotheses about PD pathophysiology and for exploring neuromodulatory interventions in a preclinical species.

## 5 Methods

### 5.1 Whole-brain simulation framework

Whole-brain activity was simulated using *The Virtual Mouse Brain* (TVMB) [74], an extension of *The Virtual Brain* (TVB) [31, 32, 75] adapted to rodent-scale resolution. The required connectivity matrices (weights *W_ij_* and delays *D_ij_*), derived from the Allen Mouse Brain Atlas [26], were obtained from standard TVMB files at 100 *µm* resolution with 596 regions (298 per hemisphere), containing log-transformed weights with soft thresholding (*w’_ij_* = *w_ij_/* min(*w*) for *w_ij_ >* 0), 99th percentile normalization, and zero diagonal, as detailed in the Allen Mouse Brain Atlas processing pipeline [26].

To reduce network complexity without sacrificing explanatory power for the dynamics of interest, regions not expected to play a primary role in Parkinsonian circuit dynamics were merged into their respective major structural groups per hemisphere, with connection strengths summed and tract lengths recomputed as connectivity-weighted averages [33]. Dynamically relevant subcortical structures (the subthalamic nucleus (STN), substantia nigra pars reticulata (SNr), ansiform lobule, cerebellar nuclei, inferior olivary complex, and specific relay thalamic nuclei) were retained as distinct nodes per hemisphere given their established roles in BG-cerebellar circuit dynamics. Each specific thalamic relay nucleus was connected unilaterally to its corresponding isocortical target, forming explicit thalamocortical loops, while self-connections were set to zero. This procedure yielded a final connectome of 212 nodes (Fig. 8).

**Fig. 8:**
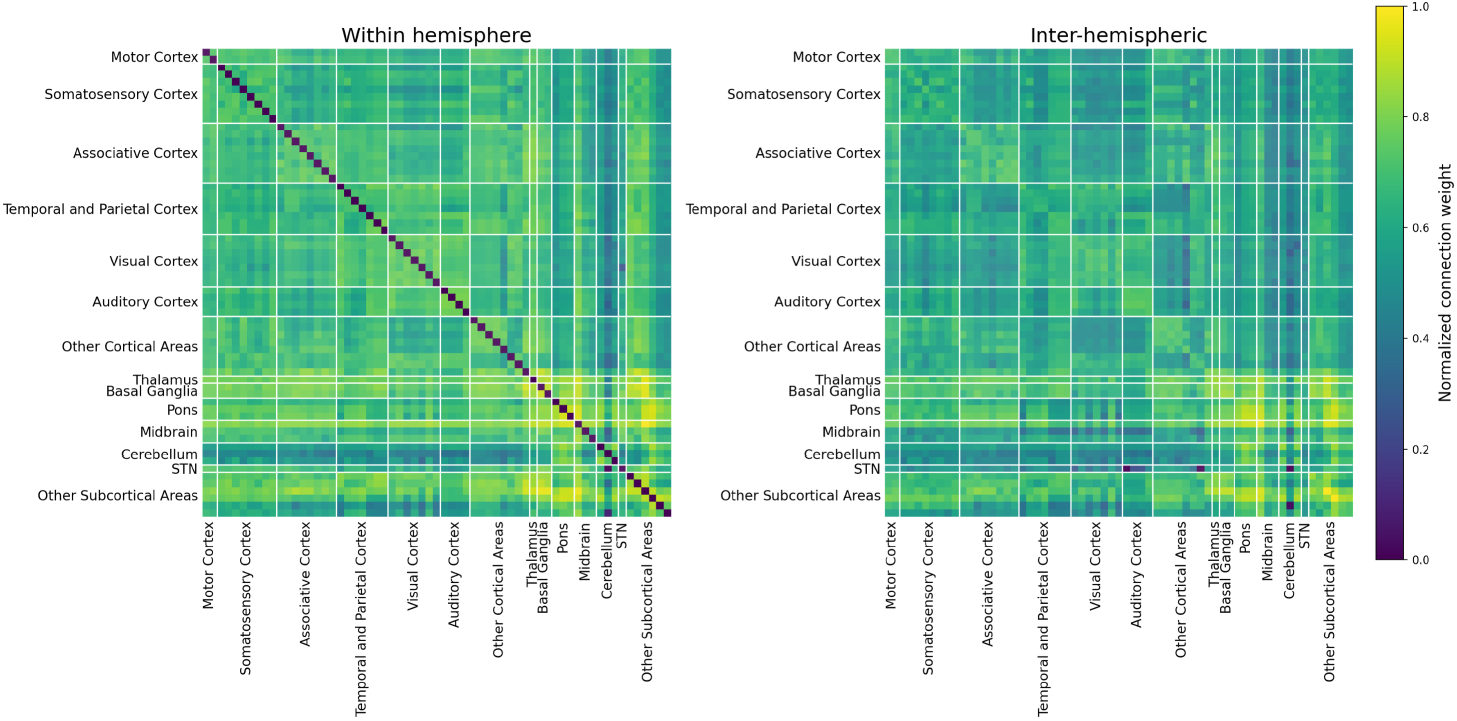
Structural connectivity used for whole-brain simulation, derived from the Allen Mouse Brain Atlas. The matrix comprises 212 regions per hemisphere, including cortical and subcortical structures.

Conduction delays were computed as:

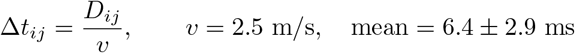

with *W_ij_∈* [0, 1] after normalisation.

### 5.2 Corticothalamic node dynamics

Each macroscopic region was modeled with the WilsonCowanThalamoCortical neural mass model within the TVB framework [33], an extension of the corticothalamic Wilson–Cowan formulation by Griffiths et al. [35], whose parameter set were slightly adapted to match mouse-specific dynamics and stable integration within the multiscale TVB–NEST framework [33]. The state *x*(*t*) of region *j* is described by three variables

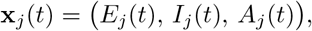

where *E_j_* and *I_j_* denote the activities of an excitatory and an inhibitory population, respectively, and *A_j_* is an auxiliary driver variable used to store the total input to the excitatory population before division by the time constant. For non-thalamic regions (cortex and subcortical nuclei), *E_j_* and *I_j_* correspond to cortical excitatory and inhibitory populations. For thalamic nuclei, *E_j_* and *I_j_* correspond to thalamic relay and reticular populations, respectively. Node identities are specified by Boolean masks *is cortical* and *is thalamic*, which determine whether a given region uses cortical or thalamic update rules. Each thalamic region connects specifically to its corresponding cortical region to form closed corticothalamic loops and receives subcortical inputs (*c_j_^sb^*) from all other subcortical regions.

All populations share the same sigmoidal activation function

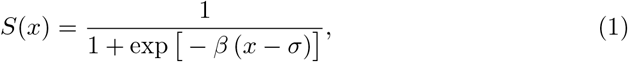

with gain *β* = 20.0 and threshold *σ* = 0.0.

#### Non-thalamic (cortical and subcortical) nodes

For regions flagged as non-thalamic (*¬is thalamic*), the dynamics of the excitatory and inhibitory populations are

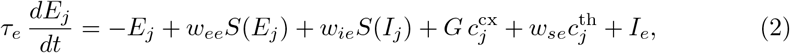

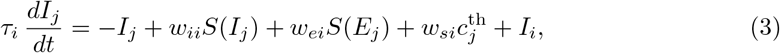

where *c_j_^cx^*, and *c_j_^sb^* and *c_j_^th^* denote long-range inputs arriving from cortical, subcortical, and thalamic relay populations, respectively (after structural connectivity, delays, and global scaling have been applied by the TVB coupling term). Subcortical input *c_j_^sb^* contributes to the excitatory drive via the same global scaling factor *G* (included in *G c_j_^cx^* in the implementation). Non-thalamic subcortical nodes lack the local thalamic feedback loop present in cortical nodes, making their dynamics more dependent on long-range connectivity and SNN-embedded regions. The auxiliary variable is updated as

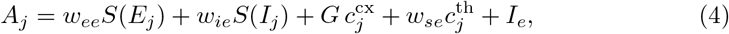

and then used to compute *dE_j_/dt* = (*−E_j_*+ *A_j_*)*/τ_e_* at each integration step (parameters summarized in Table S2 in Supplementary information.).

#### Thalamic nodes

For thalamic relay and reticular nuclei (*is thalamic*), the excitatory variable *E_j_* represents relay activity, and the inhibitory variable *I_j_* represents reticular activity. Their dynamics are

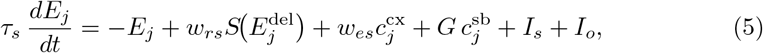

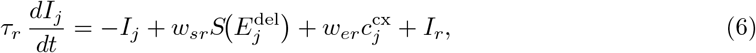

where *E_j_^del^* and *R_j_^del^* are delayed relay and reticular activities implemented via FIFO buffers, capturing local intrathalamic delays. The corresponding parameters are summarized in Table S2 in Supplementary information. Here, *τ_ct_* controls cortico–thalamic delays, and *τ_tt_* the reciprocal relay–reticular delay. The same global factor *G* scales thalamic sensitivity to subcortical input *c_j_^sb^*.

### 5.3 TVB–NEST interface architecture

The basal ganglia and cerebellar spiking networks were embedded into the whole-brain corticothalamic model using the TVB multiscale co-simulation framework [18] adapted to the rodent brain [33]. The interface provides a bidirectional mapping between rate-based neural mass variables and spiking activity, enabling stable co-evolution of the hybrid system, as follows:

**TVB → NEST:** Large-scale coupling *C_i_*(*t*) = *G*Ʃ*_j_ w_ij_S E_j_*(*t − τ_ij_*) is transformed into firing rates 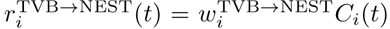 and delivered as Poisson spike trains to specific NEST populations [33]. Population-specific scaling factors *γ* were tuned via grid search to preserve standalone SNN firing statistics (firing rates in task condition, details in Subsection 5.4) when embedded in the multiscale loop. Final values:

| SNN target population | Scaling factor $\gamma$ |
| --- | --- |
| Fast-spiking interneurons (FSN) | 0.62 |
| D1/D2 medium spiny neurons | 0.55 / 0.48 |
| Subthalamic nucleus (STN) | 0.73 |
| Pontine → Granule cells | 0.90 |

**NEST → TVB:** Spiking output is first converted to mean population rates 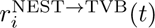, then mapped to TVB activity via low-pass filtered dynamics:

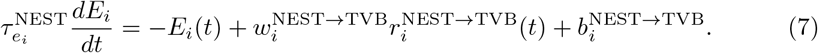

Weights were set as 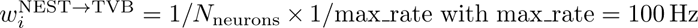 with max rate = 100 Hz, ensuring *r ∈* [0, 1] maps to TVB’s Wilson–Cowan range. For pathological high-rate regimes, an inverse-sigmoidal transformer prevents divergence:

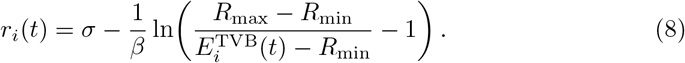

**Region-to-population mapping:** The explicit mapping is shown in Table 1.

The ansiform lobule was selected as the representative cerebellar cortical region due to its homology with primate crus I/II, implicated in tremor generation and sensorimotor coordination [76]. In mouse PD models (6-OHDA, MPTP (1-methyl-4-phenyl-1,2,3,6-tetrahydropyridine)), it shows dopamine-dependent Purkinje cell alterations contributing to beta-band tremor-like oscillations [38]. Its pontine mossy-fiber inputs from sensorimotor thalamus ensure realistic representation of cerebellum-BG interactions driving Parkinsonian tremor.

### 5.4 Basal ganglia and cerebellar spiking networks

Subcortical microcircuits were simulated in NEST 3.7 [77, 78] within the TVB-multiscale framework [18, 33], using the basal ganglia and cerebellar spiking neural networks previously adapted and validated in the multiscale model of Gambosi et al. [19]. The present implementation preserves the same neuron models, network architecture, and dopamine-dependent mechanisms as the validated standalone versions, while the complete code and parameter files provide the implementation details.

#### Basal ganglia network

The BG spiking network comprises fast-spiking interneurons (FSN), two medium spiny neuron populations (MSN D1, MSN D2; corresponding to the direct and indirect pathways), globus pallidus externa (GPe TA, GPe TI), subthalamic nucleus (STN), and substantia nigra pars reticulata (SN). These populations are instantiated bilaterally and mapped to the corresponding Allen regions as *Right/Left Striatum* (FS, MSN1, MSN2), *Right/Left Pallidum* (GPe TA, GPe TI), *Right/Left Hypothalamus* (STN), and *Right/Left Substantia nigra, reticular part* (SN) (see Table 1). The total number of neurons is distributed across populations according to the proportions inherited from the previously validated model [19], with population sizes and biophysical parameters specified in the accompanying parameter set.

Single neurons are implemented using the custom NEST models ported from the library of Lindahl et al. [36]. Striatal FSN and MSN populations follow adaptive quadratic integrate-and-fire dynamics, while GPe TA, GPe TI, STN, and SN populations are modeled as adaptive exponential integrate-and-fire units, matching the standalone BG model used in the original multiscale framework [19]. To capture biological heterogeneity, membrane and threshold parameters are drawn from distributions centered on the nominal values specified in the model configuration.

Intra-BG connectivity reproduces the architecture of the validated standalone network [19, 36]. Connection types, synapse templates, and fan-in values are inherited from the original model, and the network is instantiated using fixed-indegree rules with weights drawn around the nominal values. For FS–MSN and MSN–MSN projections, connectivity is spatially restricted to preserve the local organization of the striatum. Dopamine depletion in the BG is controlled by a scalar parameter *ξ ∈* [−0.8, 0] passed from the global configuration and applied in the same way as in Gambosi et al. [19], so that dopamine-dependent changes in excitability and synaptic gain are inherited from the previously validated model. In standalone mode, the BG network can also receive tonic Poisson drive, but in the multiscale configuration this drive is replaced by TVB-derived input through the co-simulation interface. Spiking activity is recorded and converted into firing-rate traces for coupling to the corticothalamic mass model.

#### Cerebellar network

The cerebellar spiking network follows the EGLIF-based olivocerebellar microcircuit of Geminiani et al. [37], as adapted in our previous multiscale work [19]. The network includes the major cerebellar populations: glomeruli (Glom), granule cells (GrC), Golgi cells (GoC), Purkinje cells (PC), basket and stellate cells (BC, SC), deep cerebellar nuclei neuron types (DCN and DCNp), and inferior olive (IO). These populations are instantiated bilaterally and mapped to the corresponding Allen regions as *Right/Left Ansiform lobule* (granule, Golgi, basket, stellate, Purkinje, glomeruli), *Right/Left Cerebellar nuclei* (DCN, DCNp), and *Right/Left Inferior olivary complex* (IO) (see Table 1). The neuron and synapse models are implemented in the NEST module CerebNEST. Their parameters follow the validated cerebellar implementation, and the full parameterization is available in the shared code.

The cerebellar connectivity is reconstructed from the 3D topology of the original model [79], preserving the microcomplex organization and the main mossy-fiber and climbing-fiber projections. Synaptic weights, delays, and receptor identities are inherited from the validated network description. Dopamine depletion is implemented through the same Purkinje-cell loss mechanism used in the original model, with the surviving fraction varying as a function of the depletion level to reproduce the pathological reduction in PC number reported in mouse lesion data [19, 38]. The surviving fraction depends linearly on the depletion level:

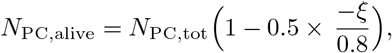

where *ξ ∈* [0, −0.8] is shared with the BG model. In standalone simulations, low-rate Poisson input can be applied to glomeruli and task-like sensory drive can be provided through mossy-fiber pathways, but in the multiscale configuration these external sources are disabled and replaced by TVB-derived input. As for the BG network, spiking output is recorded and transformed into firing-rate signals for coupling to the whole-brain model.

### 5.5 Simulation protocol

Two conditions were simulated: a *physiological* baseline and a set of *Parkinsonian* states generated by applying dopamine depletion within the spiking basal ganglia (BG) and cerebellar networks using *ξ ∈ {−*0.1, −0.2, −0.4, −0.8*}*. Each simulation ran for 3500 ms, with the initial 25% discarded to remove transients. Independent noise realisations were used for all trials.

### 5.6 Spectral analysis

Oscillatory activity was quantified from regional time series using a continuous wavelet transform (CWT) to estimate power across canonical frequency bands for mice [80, 81] (delta: 0.5–4 Hz; theta: 4–10 Hz; beta: 10–30 Hz; gamma: 30–100 Hz). Relative changes were computed as:

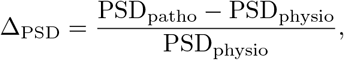

and group differences were evaluated using Wilcoxon–Mann–Whitney tests with FDR correction.

Then, high-resolution analysis employed a Morlet CWT (1–60 Hz, 240 logarithmic frequencies, *w* = 10) followed by “*Fitting Oscillations and One-Over F*” FOOOF-based decomposition (aperiodic mode=fixed) [82]. Only the periodic component was integrated within canonical bands, yielding aperiodic-corrected oscillatory power. Both absolute and relative changes were computed per trial.

### 5.7 Regional grouping for systems-level analysis

To facilitate interpretation at the system level, the 212 regions of the parcellation were aggregated into anatomically and functionally coherent macro-domains derived from the Allen Mouse Brain Atlas. Table 2 lists all groups and their corresponding region indices.

**Table 2:**
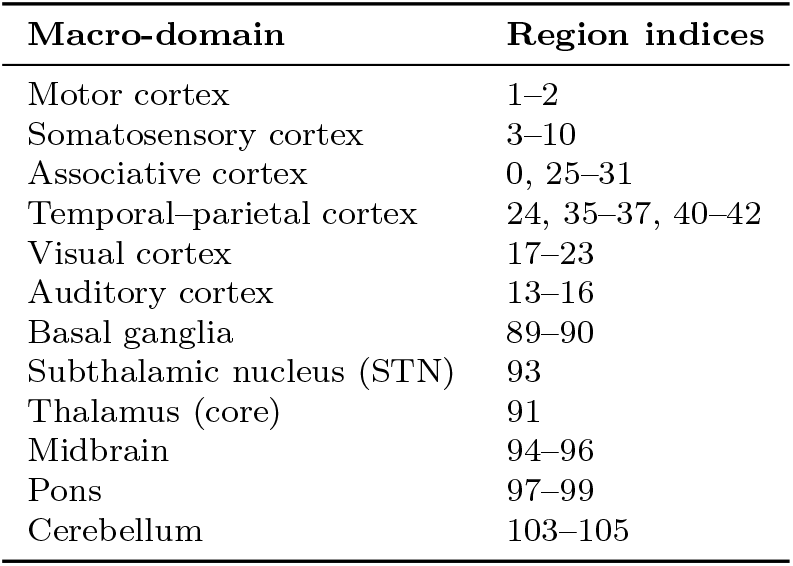
Regional macro-domains used for spectral analyses.

| Macro-domain | Region indices |
| --- | --- |
| Motor cortex | 1–2 |
| Somatosensory cortex | 3–10 |
| Associative cortex | 0, 25–31 |
| Temporal–parietal cortex | 24, 35–37, 40–42 |
| Visual cortex | 17–23 |
| Auditory cortex | 13–16 |
| Basal ganglia | 89–90 |
| Subthalamic nucleus (STN) | 93 |
| Thalamus (core) | 91 |
| Midbrain | 94–96 |
| Pons | 97–99 |
| Cerebellum | 103–105 |

Two analyses were conducted:

1. **Dopamine-progression analysis (beta-specific).** We quantified how *β*-band (10–30 Hz) power scaled with dopamine depletion across macro-domains using *ξ* = *{*0, −0.1, −0.2, −0.4, −0.8*}*.
2. **Full-band analysis in severe depletion.** In the most pathological state (*ξ* = −0.8), we evaluated spectral power across all bands (*δ*, *θ*, *β*, *γ*) to characterise global reorganisation beyond the *β* range.

This grouping strategy enables a compact systems-level description of how oscillatory abnormalities propagate across cortical, subcortical and cerebellar circuits.

### 5.8 In-silico ablation experiment: isolating resonance mechanisms

To test whether *β* oscillations arise intrinsically within BG circuits or require cortico– subcortical resonance, we performed a targeted perturbation experiment in which all long-range inputs from cortex and thalamus to the BG SNN were removed. This was implemented by setting the TVB→BG interface gains to zero:

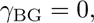

for all D1, D2, FS, and STN input channels, while preserving internal BG connectivity, BG→TVB output projections, and the entire cerebellar pathway.

The hybrid model was then simulated under the same protocol as above, and spectral power was estimated using the identical CWT pipeline. Power was computed across EEG bands adapted for mice [80]:

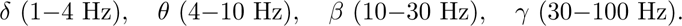

Comparison with the intact network enabled isolation of intrinsic BG dynamics from loop-mediated amplification, providing a mechanistic test of resonance hypotheses at the whole-brain scale.

## Supplementary information

**Fig. S1**:Validation of population firing rates in the cerebellar and basal ganglia modules. Mean firing rates for the main neuronal populations in the cerebellar network (top) and basal ganglia network (bottom) are compared between the values reported in Gambosi et al. (2024) and those obtained in the present TVB multiscale model. **Table S2**: Parameters of the corticothalamic Wilson-Cowan model. **Table S3**: List of abbreviations.

## Acknowledgements

The authors acknowledge the following fundings: BG, AA, AP acknowledges support by SGA 945539 (Human Brain Project SGA3) and Horizon Europe Programme for Research and Innovation, Grant Agreement No. 101147319 (EBRAINS 2.0); JM acknowledges support by the Deutsche Forschungsgemeinschaft (DFG, German Research Foundation) - Project-ID 424778381 - TRR 295; PR acknowledges supported by the EU Horizon Europe program: BRIDGE (101219311), EBRAINS2.0(101147319), Virtual Brain Twin (101137289), EBRAINS-PREP 101079717, AISN 101057655, EBRAIN-Health 101058516, EIC grant PHRASE 101058240, the Digital Europe Programme: TEF-Health (101100700), SHAIPED (101195135), CoordinaTEF (101168074), the German Research Foundation: SFB 1436 (project ID 425899996); SFB 1315 (project ID 327654276); SFB 936 (project ID 178316478; SPP Computational Connectomics RI 2073/6-1, RI 2073/10-2, RI 2073/9-1; DFG Clinical Research Group BECAUSE-Y 504745852, DFG German-Spanish project BrainsAge RI 2073/14-1, DFG Project-ID 424778381 - TRR 295 Retune, ERA PerMed JTC2021 German Federal Ministry for Health PatternCog ZMI5-2522FSB904,Berlin University Alliance OpenMake, the Virtual Research Environment at the Charité Berlin and EBRAINS Health Data Cloud and the Berlin Institute of Health and Foundation Charité.

## Declarations

The authors declare no competing interests.

## Ethics approval and consent to participate

Not applicable. This study uses only computational simulations.

## Code and data availability

Simulation code and derived data are available on github at this repository. Publicly available connectivity data were obtained from the Allen Mouse Brain Atlas [26].

---

**Fig. S1:**
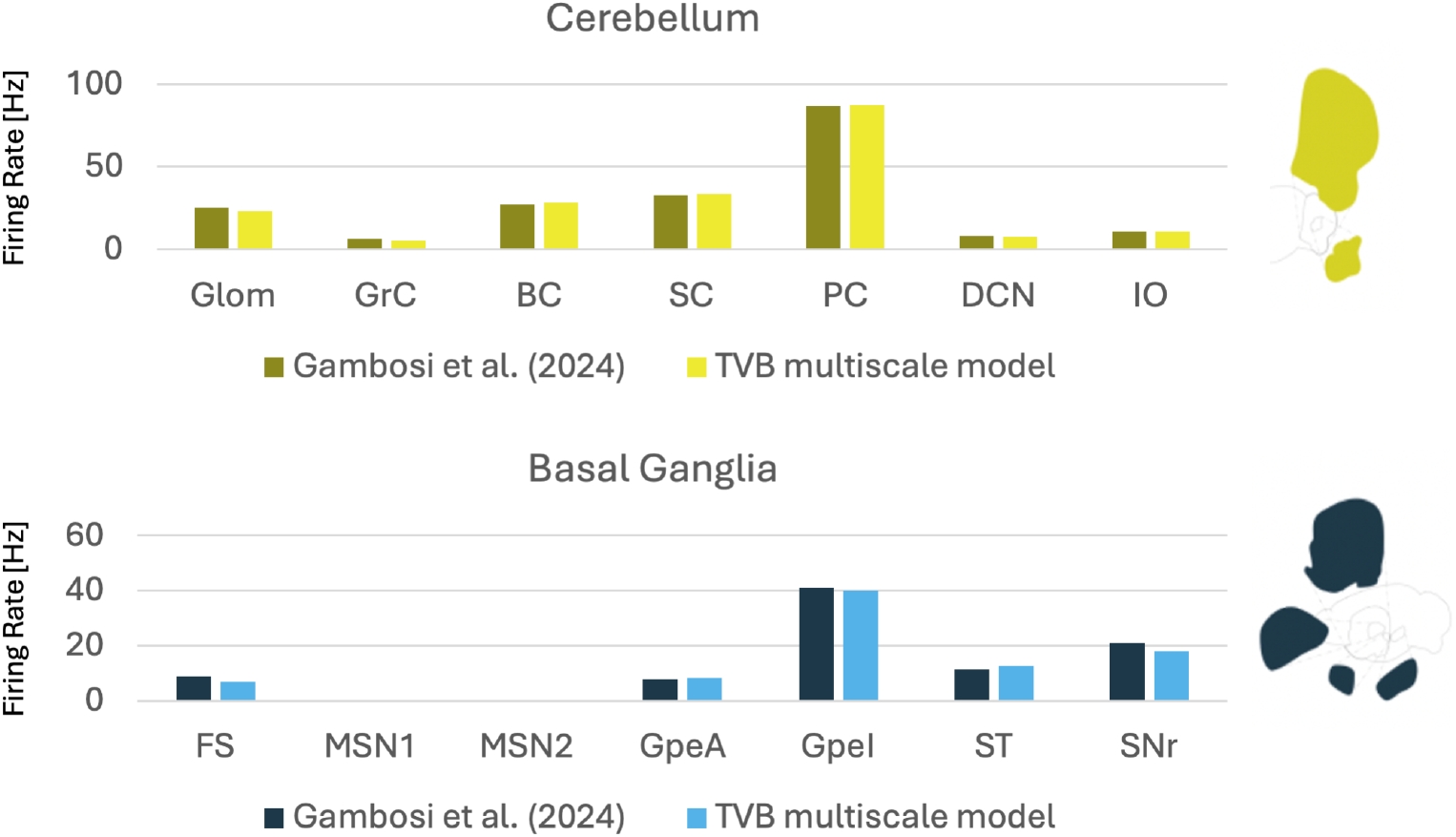
Validation of population firing rates in the cerebellar and basal ganglia modules. Mean firing rates for the main neuronal populations in the cerebellar network (top) and basal ganglia network (bottom) are compared between the values reported in Gambosi et al. (2024) and those obtained in the present TVB multiscale model.

**Table S2:**
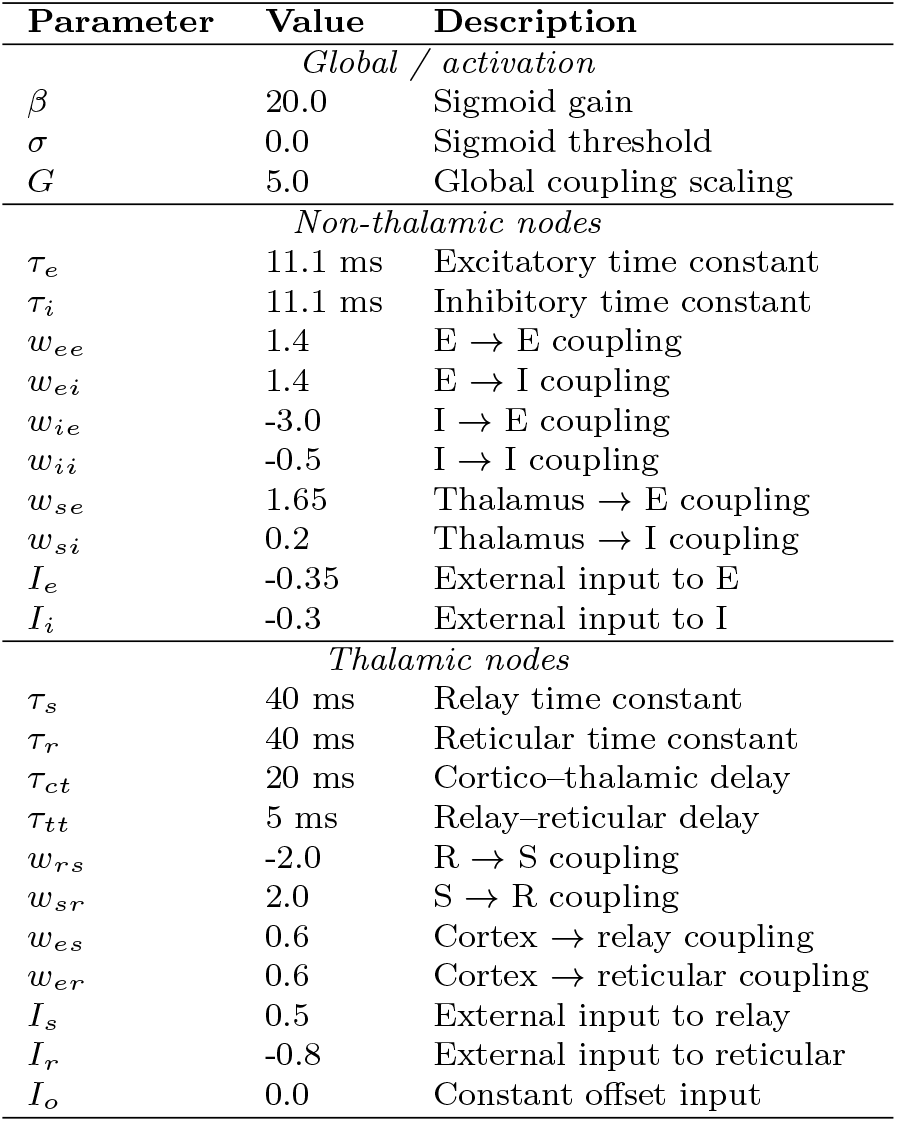
Parameters of the corticothalamic Wilson–Cowan model.

**Table S3:**
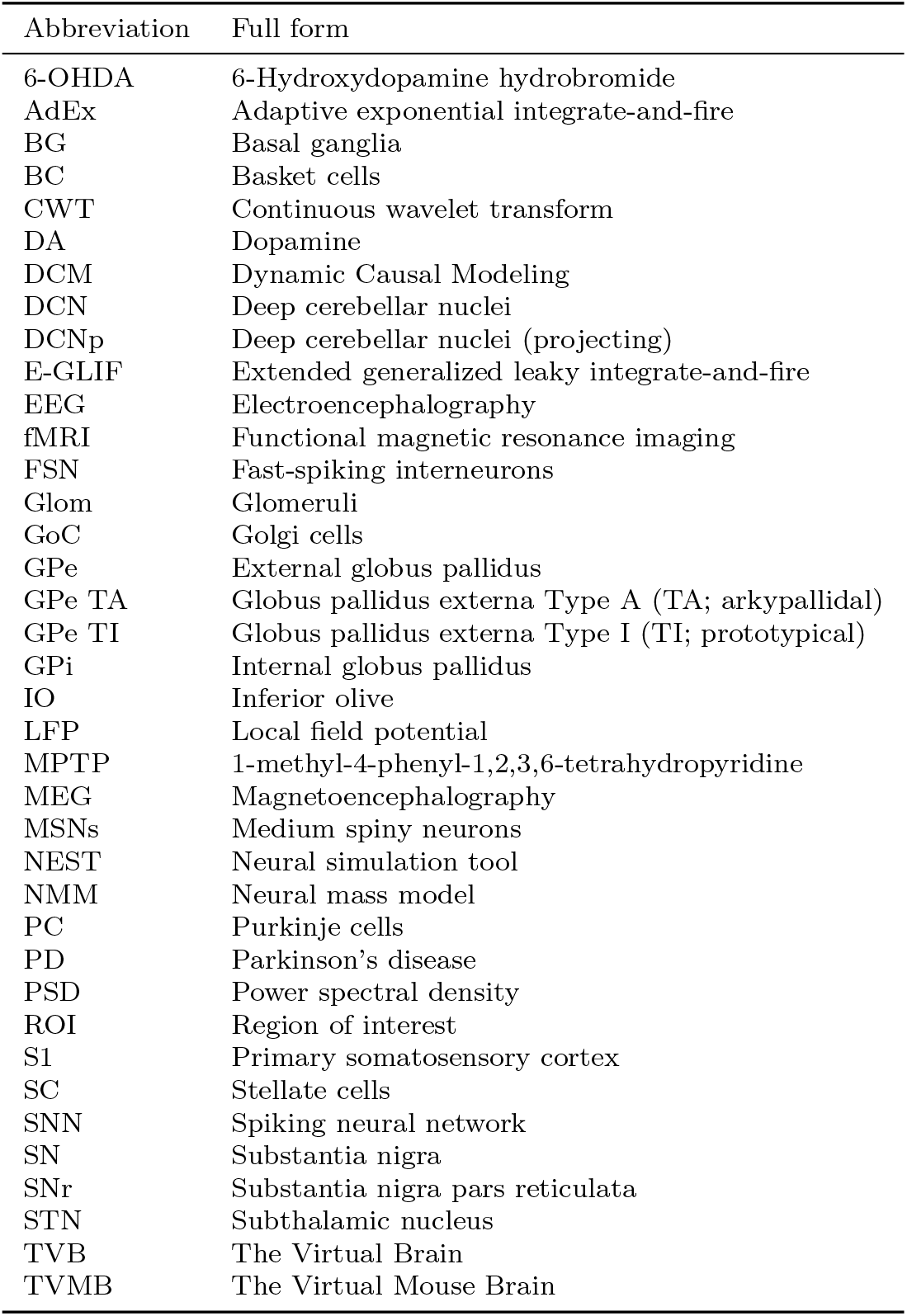
List of abbreviations.

