## Supplementary for "Dopamine Depletion Drives Whole-Brain Oscillatory Disruptions via Cortico–Subcortical Resonance: A Multiscale Model of Parkinson’s Disease in Mice"

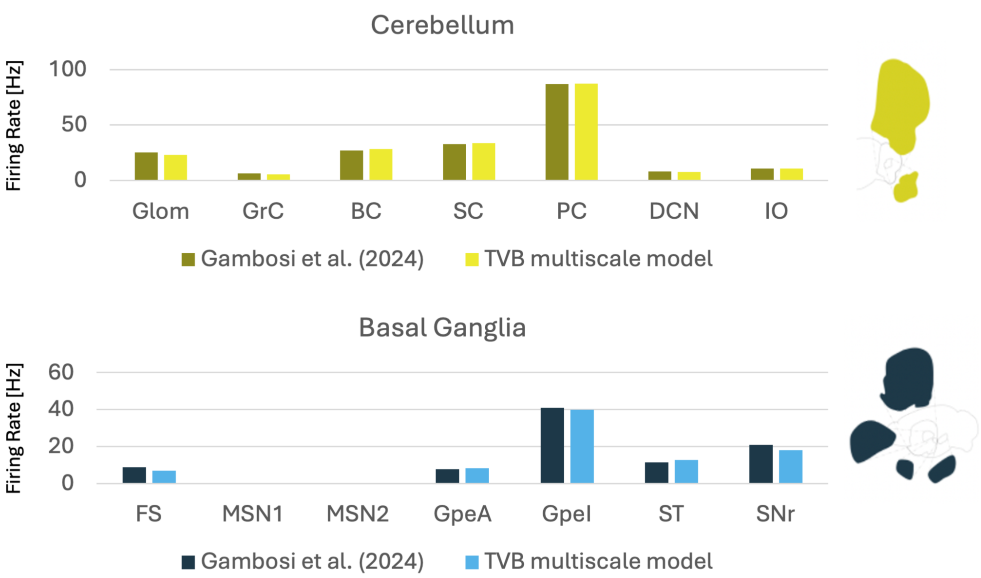


**Fig. S1**: Validation of population firing rates in the cerebellar and basal ganglia modules. Mean firing rates for the main neuronal populations in the cerebellar network (top) and basal ganglia network (bottom) are compared between the values reported in Gambosi et al. (2024) and those obtained in the present TVB multiscale model.

**Table S2**: Parameters of the corticothalamic Wilson–Cowan model.

| Parameter | Value | Description |
|---|---|---|
| | *Global / activation* | |
| $\beta$ | 20.0 | Sigmoid gain |
| $\sigma$ | 0.0 | Sigmoid threshold |
| $G$ | 5.0 | Global coupling scaling |
| | *Non-thalamic nodes* | |
| $\tau_e$ | 11.1 ms | Excitatory time constant |
| $\tau_i$ | 11.1 ms | Inhibitory time constant |
| $w_{ee}$ | 1.4 | E $\rightarrow$ E coupling |
| $w_{ei}$ | 1.4 | E $\rightarrow$ I coupling |
| $w_{ie}$ | -3.0 | I $\rightarrow$ E coupling |
| $w_{ii}$ | -0.5 | I $\rightarrow$ I coupling |
| $w_{se}$ | 1.65 | Thalamus $\rightarrow$ E coupling |
| $w_{si}$ | 0.2 | Thalamus $\rightarrow$ I coupling |
| $I_e$ | -0.35 | External input to E |
| $I_i$ | -0.3 | External input to I |
| | *Thalamic nodes* | |
| $\tau_s$ | 40 ms | Relay time constant |
| $\tau_r$ | 40 ms | Reticular time constant |
| $\tau_{ct}$ | 20 ms | Cortico–thalamic delay |
| $\tau_{tt}$ | 5 ms | Relay–reticular delay |
| $w_{rs}$ | -2.0 | R $\rightarrow$ S coupling |
| $w_{sr}$ | 2.0 | S $\rightarrow$ R coupling |
| $w_{es}$ | 0.6 | Cortex $\rightarrow$ relay coupling |
| $w_{er}$ | 0.6 | Cortex $\rightarrow$ reticular coupling |
| $I_s$ | 0.5 | External input to relay |
| $I_r$ | -0.8 | External input to reticular |
| $I_o$ | 0.0 | Constant offset input |

**Table S3**: List of abbreviations

| Abbreviation | Full form |
|---|---|
| 6-OHDA | 6-Hydroxydopamine hydrobromide |
| AdEx | Adaptive exponential integrate-and-fire |
| BG | Basal ganglia |
| BC | Basket cells |
| CWT | Continuous wavelet transform |
| DA | Dopamine |
| DCM | Dynamic Causal Modeling |
| DCN | Deep cerebellar nuclei |
| DCNp | Deep cerebellar nuclei (projecting) |
| E-GLIF | Extended generalized leaky integrate-and-fire |
| EEG | Electroencephalography |
| fMRI | Functional magnetic resonance imaging |
| FSN | Fast-spiking interneurons |
| Glom | Glomeruli |
| GoC | Golgi cells |
| GPe | External globus pallidus |
| GPe TA | Globus pallidus externa Type A (TA; arkypallidal) |
| GPe TI | Globus pallidus externa Type I (TI; prototypical) |
| GPi | Internal globus pallidus |
| IO | Inferior olive |
| LFP | Local field potential |
| MPTP | 1-methyl-4-phenyl-1,2,3,6-tetrahydropyridine |
| MEG | Magnetoencephalography |
| MSNs | Medium spiny neurons |
| NEST | Neural simulation tool |
| NMM | Neural mass model |
| PC | Purkinje cells |
| PD | Parkinson's disease |
| PSD | Power spectral density |
| ROI | Region of interest |
| S1 | Primary somatosensory cortex |
| SC | Stellate cells |
| SNN | Spiking neural network |
| SN | Substantia nigra |
| SNr | Substantia nigra pars reticulata |
| STN | Subthalamic nucleus |
| TVB | The Virtual Brain |
| TVMB | The Virtual Mouse Brain |
